# DUSP4 knockdown in BRAF V600E mutant colorectal cancer induces cell cycle arrest and halts tumor growth

**DOI:** 10.64898/2026.01.15.699635

**Authors:** Barbara Hopfgartner, Monika Kriz, Donat Alpar, Yvonne Westermaier, Marco H. Hofmann, Ralph A. Neumüller, Sarah Grosche

## Abstract

Oncogenic signaling in cancer cells is essential for proliferation, and its disruption, either through inhibition or overactivation, can provide therapeutic opportunities. Dual specificity phosphatase 4 (DUSP4), a negative regulator of the MAPK pathway that dephosphorylates ERK, has been proposed as a potential target; however, its therapeutic relevance has not been evaluated *in vivo*. In this study, we show that DUSP4 knock-down induces G1 cell cycle arrest and reduces proliferation in BRAF^V600E^-mutant and BRAF inhibitor-resistant colorectal cancer models, both *in vitro* and *in vivo,* and induces a rapid DUSP5-mediated adaptive response. While treatment achieved tumor stasis indicating disease control, it did not yield tumor regression, suggesting that DUSP4 may have limited efficacy as a monotherapy target in cancer.

## Introduction

The MAPK pathway is tightly regulated in both normal tissues and pathological contexts. More than 40% of human cancers involve alterations in this pathway^1^. These alterations include activating mutations and/or amplifications at the level of upstream receptors (e.g., EGFR, HER2) or at various nodes (SOS1, KRAS, NRAS, HRAS, BRAF) in the signaling cascade^2–5^. BRAF^V600E^ is a well-characterized oncogenic mutation in the MAPK pathway, present in approximately 50% of melanomas and 10% of colorectal cancers^6,7^.

The consequences of enhanced oncogenic signaling depend on context and signaling strength. In melanocytes, BRAF^V600E^ expression is associated with oncogene-induced senescence^8,9^. In contrast, BRAF^V600E^ mutation-positive melanoma cells remain dependent on sustained signaling, as demonstrated by tumor regressions following BRAF inhibitor treatment^10–12^.

While inhibition of oncogenic signaling has been a successful therapeutic strategy, recent studies suggest that overactivation of oncogenic signaling could represent a novel therapeutic approach^13–17^. Dual specificity phosphatase 4 (DUSP4), a negative regulator of the MAPK pathway through dephosphorylation of ERK, has been proposed as a potential therapeutic target^17,18^. However, its relevance for *in vivo* tumor growth remains unclear. Our data indicate a measurable but limited impact on tumor growth and a rapid adaptive response, providing valuable insights into the biology of DUSP4 and casting doubt on the viability of DUSP4 as a monotherapy target.

## Methods

### Cell lines

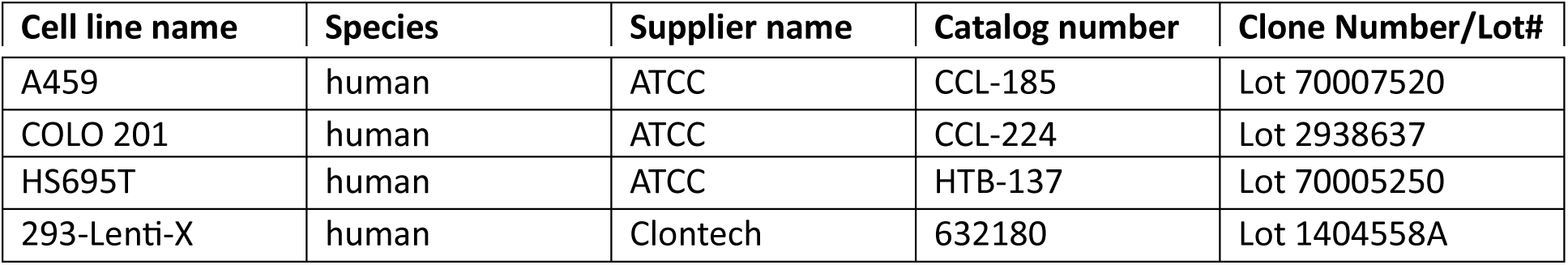

### Antibodies

Anti-DUSP4 (Cell Signaling, Cat. No. 5149); anti-α-Actinin (Cell Signaling, Cat. No. 3134); anti-GAPDH (Abcam, Cat. No. Ab181602); DUSP1/MKP1 (Cell Signaling, Cat. No. 35217); anti-DUSP5 Abcam, Cat. No. Ab200708).

### Sequences

All gRNAs are listed in Supplementary Table 1.

### Compounds

Trametinib was purchased from Selleckchem.com (Cat. No. S2673; Batch No. 8). Dabrafenib Tafinlar®, a potent ATP-competitive inhibitor of b-Raf, b-Raf(V600E) and c-Raf from Novartis Pharma GmbH, was used in *in vitro* experiments (CAS 1195765-45-7). Vemurafenib Zelboraf®, a potent ATP-competitive inhibitor of b-Raf, b-Raf(V600E) and c-Raf from Plexxikon/Roche, was used in *in vitro* experiments (CAS 918504-65-1). Doxycycline hyclate was purchased from Sigma Aldrich in *in vivo* experiments (Cat. No. D9891) and in *in vitro* experiments (Cat. No. D3447).

### Competitive Proliferation Assay

A lentiviral expression vector encoding Cas9 and puromycin was stably integrated into the target cells. Selected cells were transduced with a lentivirus carrying sgRNA expression vectors at an infection efficiency of 20–70%. Infection levels were determined by flow cytometry based on GFP expression three days after infection (designated as Day 0). SgRNA-positive cells were monitored at regular intervals, and all time points are shown relative to Day 0. For rescue experiments, cells were transduced with either a lentiviral tet-inducible cDNA vector or a constitutive cDNA vector. For the tet-inducible vector, cDNA expression was induced with 0.5 µg mL^−1^ doxycycline at the time of sgRNA infection, and the percentage of sgRNA-positive cells was monitored by flow cytometry at regular intervals.

### Simple Western

Cells were lysed in RIPA buffer. Samples were then mixed with Simple Western Sample Buffer and standards to a final concentration of 0.5 μg/μL and heat denatured at 95°C. The prepared samples, primary and secondary antibodies, and the chemiluminescent substrate were dispensed into designated wells according to the Simple Western protocol. The Simple Western capillary and the prepared assay plate were analyzed using ProteinSimple technology by Bio-Techne (Catalog # 004-650) which automatically carried out all assay steps.

### RT-PCR

cDNA synthesis and RT-PCR from whole-cells were performed using the FastLane Cell Multiplex Kit (Qiagen, Cat. No. 216513) and a QuantStudio™ 5 Real-Time PCR system. RNA from tumor tissue was extracted using the RNeasy MiniKit (Qiagen, Cat. No. 74106) according to the supplier’s instructions. Quantitative PCR was performed using the QuantiTect Multiplex RT-PCR Kit (Qiagen, Cat. No. 204643) and a QuantStudio™ 5 Real-Time PCR system.

The following primers were used: Applied Biosystems™ Human GAPD (GAPDH) Endogenous Control (VIC™/MGB probe, primer limited) (ThermoFisher, Cat. No. 43-263-17E). Applied Biosystems™ TaqMan™ Gene Expression Assay (FAM) (ThermoFisher, Cat. No. 4331182):

|  |  |
| --- | --- |
| **CCND1** | Hs00765553_m1 |
| **DUSP1** | Hs00610256_g1 |
| **DUSP4** | Hs01027785_m1 |
| **DUSP5** | Hs00244839_m1 |
| **DUSP6** | Hs01027785_m1 |
| **EPHA2A** | Dr03074113_m1 |
| **EPHA4** | Hs00953178_m |
| **ETV4** | Hs00383361_g1 |
| **ETV5** | Hs00927557_m |
| **PHLDA1** | Hs00378285_g1 |
| **SPRY2** | Hs01921749_s1 |
| **SPRY4** | Hs01935412_s1 |

### Cell Viability Assay

Twenty-four hours prior to treatment, cells were plated in a 96-well plate. All treatments were performed in technical duplicates. Treated cells were incubated for six days at 37°C with 5% CO_2_. The CellTiter-Glo® Luminescent Cell Viability Assay was performed, and luminescence signals were detected using a VICTOR X4 multilabel plate reader. Quantifications of viable cells were calculated by normalizing treated cells to DMSO-treated controls.

### Plasmids

For the competitive proliferation assay, sgRNAs were individually expressed from the Lenti_gRNA-GFP(LRG)_2.1T. Cas9 was expressed using the LentiV_Cas9_puro. For the knockout-rescue experiment, sgRNAs were expressed from the LRG vector, and cDNA was expressed from a tet-inducible UTR3GV-IRES-Neo vector.

To generate the COLO 201 ARTi cell line, sgRNAs were cloned into pX458, which harbors the wild-type Cas9 coding region, a gRNA expression cassette, and a GFP reporter. A pUC57-Simple expression vector containing the respective donor sequence was used as the repair template.

### Cell Culture and Media

Packaging cells (Lenti-X™ 293T Cell Line) used for producing pantropic lentiviral particles and MIA PaCa-2 were cultured in DMEM supplemented with 10% FBS at 37°C with 5% CO_2_. COLO 201 cells were cultured in RPMI supplemented with 10% FBS at 37°C with 5% CO_2_. Hs 695T cells were cultured in EMEM supplemented with 10% FBS, sodium pyruvate (1 mM), and GlutaMAX (4 mM) at 37°C with 5% CO_2_. A549 cells were cultured in Ham’s F-12K (Kaighn’s) supplemented with 10% FBS at 37°C with 5% CO_2_.

### Generation of Resistant Cell Line and RNA Sequencing

The human colorectal adenocarcinoma cell line COLO 201 Cas9 Puro, which carries a BRAF^V600E^ mutation and constitutively expresses Cas9, was treated for more than two months with 10x the IC_50_ concentration of dabrafenib until a resistance-associated IC_50_ shift was detected. After confirming the resistance stability, the remaining cell pool was validated by comparing the IC_50_ shift to that of the parental cell line maintained under identical culture conditions. Subsequently, resistant and non-resistant cells were infected with gRNAs (AAVS1 and DUSP4) and sorted five days post-infection. All conditions were performed in triplicates. Total RNA was purified from 0.5-1 M cells per sample using the RNeasy Plus Universal Kit (Qiagen). Extracted RNA samples were normalized to 100 ng/µl in 10 µl nuclease-free H_2_O. From each sample, 500 ng of total RNA was subjected to genome-wide gene expression profiling via 3’-mRNA sequencing. Next generation sequencing (NGS) libraries were prepared using the QuantSeq 3’-mRNA-Seq V2 Library Prep Kit FWD for Illumina (Lexogen GmbH, Vienna, Austria) according to the manufacturer’s protocol using the Biomek i7 automated liquid handling workstation (Beckman Coulter GmbH, Krefeld, Germany). After quality control with High Sensitivity D1000 ScreenTape on a 4200 TapeStation System (Agilent Technologies Österreich GmbH, Vienna, Austria), individual libraries were diluted, pooled equimolarly, quantified using the NEBNext Library Quant Kit for Illumina (New England BioLabs Inc., #E7630L), and sequenced on a NextSeq 2000 instrument (Illumina, San Diego, CA) using 75 bp single-read chemistry.

### ARTi Cell Line Generation

To generate COLO 201 ARTi cells, gRNAs targeting the end of the DUSP4 locus were cloned into pX458, which harbors the wild-type Cas9 coding region, a gRNA expression cassette, and a GFP reporter. The CRISPR construct and the corresponding DUSP4 Hygromycin ARTi repair template were transiently transfected into COLO 201 cells carrying rtTA and an inducible mirF cassette against the ARTi sequence, as described in Hoffmann *et al*.^19^. For transfection, 3 × 10^5^ cells were plated per well in a 6-well dish 24 h prior to transfection. Cells were transfected using FuGENE® HD Transfection Reagent. Subsequently, the cells were cultured for six days to allow genome editing. After editing, cells were placed in medium supplemented with 100 µg/mL hygromycin to select for edited cells. Afterwards, cells were plated into 96-well plates at single cell density. Growing colonies were then selected, and gDNA and protein lysates were collected for further analysis. Genome-engineered cells were identified by PCR amplification of CRISPR/Cas9-induced genomic modifications, Sanger Sequencing, and Simple Western analysis.

### RealTime-Glo™ Annexin V Apoptosis and Necrosis Assay

The RealTime-Glo™ Annexin V Apoptosis and Necrosis Assay (Promega, Cat. #JA1011) was used to monitor apoptosis and necrosis in real time. The assay was performed according to the manufacturer’s instructions. COLO 201 ARTi and COLO 201 ARTi dabrafenib-resistant cells were seeded in a 96-well plate at a density of 1 x 10⁴ cells per well in medium containing 0.5 µg/mL doxycycline to induce DUSP4 mRNA degradation. As positive control, COLO 201 ARTi and COLO 201 ARTi dabrafenib-resistant cells with no doxycycline were treated with 2.5 µM staurosporine to induce apoptosis and 10 µM raptinal to induce necrosis. The detection reagent was added directly to the wells. Luminescence and fluorescence signals were measured using a multimode plate reader (Envision) for four days. Luminescence indicated Annexin V binding to phosphatidylserine (PS) on the outer leaflet of the cell membrane, while fluorescence indicated loss of membrane integrity. The onset of apoptosis was determined by an increase in luminescence, whereas secondary necrosis was detected by an increase in fluorescence. Data were normalized to the untreated control and expressed as mean ± standard deviation (SD) from triplicates.

### Click-iT® Plus EdU Flow Cytometry Assay

The Click-iT® Plus EdU Flow Cytometry Assay (Thermo Fisher Scientific, Cat. #C10635) was used to measure cell proliferation. The assay was performed according to the manufacturer’s instructions. COLO 201 ARTi and COLO 201 ARTi dabrafenib-resistant cells were seeded in a 24-well plate at a density of 4 x 10^4^ cells per well. Cells were treated with 0.5 µg/mL of doxycycline for 24 to 96 h. As a positive control, cells were treated with 0.5 µM or 1 µM nocodazole for 24 h. EdU was added to the culture medium at a final concentration of 10 µM, and cells were incubated for 2 h at 37°C. After incubation, cells were harvested by centrifugation at 1500 rpm for 5 min and washed twice with 1% bovine serum albumin (BSA) in phosphate-buffered saline (PBS). Cells were fixed and permeabilized with 100 µL Click-iT® fixative for 15 min at room temperature, protected from light. Cells were then washed twice with 1% BSA in PBS. After the second wash, the cell pellets were resuspended in 1X Click-iT® saponin-based permeabilization and wash reagent and incubated for 15 min. The Click-iT® reaction cocktail was added to the cells and incubated for 30 min at room temperature, protected from light. Afterwards, cells were washed twice with 1X Click-iT® saponin-based permeabilization and wash reagent. The supernatant was removed, and the pellet was resuspended in 100-200 µL of complete media. Cells were additionally stained with Vybrant^TM^ DyeCycle^TM^ Violet stain (Thermo Fisher Scientific, Cat. #V35003) at a final concentration of 5 µM and incubated at 37°C for 30 min, protected from light. All conditions were performed in triplicates. A Beckman Coulter CytoFLEX was used to analyze the samples.

### Dabrafenib-Resistant Cell Line Generation (HS695T and COLO 201 DUSP4 ARTi)

*In vitro* induction of resistance in HS695T, COLO 201, and COLO 201 DUSP4 ARTi cells was achieved by exposing them to 10x their respective IC_50_ concentrations of dabrafenib over a period of two months. For HS695T, 400 nM dabrafenib was used, and for COLO 201, 30 nM of dabrafenib as applied until a detectable IC_50_ shift indicating resistance was observed. Subsequently, cells were cultured without dabrafenib for a further three weeks to ensure stability of the resistant phenotype. Stability was confirmed by verifying the IC_50_ shift compared to the parental cell after off-treatment of dabrafenib.

### Cell Line-Derived Efficacy Studies and Biomarker Studies in Mice

All animal studies were approved by the internal ethics committee and the local Austrian governmental authority. Group sizes for efficacy studies were determined based on power analysis. Female BomTac:NMRI-*Foxn1*^nu^ mice were used for both efficacy and biomarker studies. Mice were subcutaneously engrafted with 5 x 10^6^ COLO 201 ARTi cells suspended in 100 µl of 1xPBS containing 5% FCS and 50% Matrigel. For the efficacy study, tumors were randomized by size by using the automated storage data system Sepia Noto into groups of 7 mice once tumors reached a volume of 150.3 - 237.9 mm³. Mice in the control group received drinking water supplemented with 5% saccharose *(ad libitum)*, while mice in the treatment group received water containing 2mg/kg doxycycline + 5% saccharose for 29 days. Tumor size was measured twice weekly using an electronic caliper, and body weight was monitored daily. The analysis largely followed the method described in this reference^20^. On the final day of the efficacy study (day 29), tumors were excised and snap-frozen in liquid nitrogen for biomarker analysis.

### Bioinformatics Analyses

#### CRISPR Dependency Data

CRISPR screen dependency data (Chronos scores), cell line expression data, and cell line metadata (version 22Q2) were downloaded from the DepMap Consortium portal (https://depmap.org/portal). Heatmaps were generated using R version 4.4.0 with the following packages: ComplexHeatmap (version 2.20.0), data.table_1.16.0, ggplot2_3.5.1, circlize_0.4.16, tidyr_1.3.1, RColorBrewer_1.1-3, and dplyr_1.1.4. Column clustering was performed using the default settings of the ComplexHeatmap package.

### RNA Sequencing Analysis

Single-end sequencing reads from grafted samples were filtered into human and mouse reads using Disambiguate^21^, based on mapping to hg38 and mm10. The filtered reads were then processed using a pipeline that builds upon and extends the ENCODE “Long RNA-seq” pipeline: Filtered reads were aligned to either *Homo sapiens* (human) genome hg38/GRCh38 (primary assembly, excluding alternate contigs) or the *Mus musculus* (mouse) genome mm10/GRCm38 using the STAR aligner (v2.5.2b)^22^ allowing for soft clipping of adapter sequences. For quantification, we used transcript annotation files from Ensembl version 86, corresponding to GENCODE 25 for human and GENCODE M11 for mouse. Samples were quantified using these annotations with RSEM (v1.3.0)^23^ and featureCounts (v1.5.1)^24^. Quality controls was performed at the respective steps using FastQC (v0.11.5) (Andrews S. 2010; available at: http://www.bioinformatics.bbsrc.ac.uk/projects/fastqc), FastQC (a quality control tool for high-throughput sequence data (http://www.bioinformatics.babraham.ac.uk/projects/fastqc)), picardmetrics (v0.2.4) (Slowikowski K. (2016): Available online at: https://github.com/slowkow/picardmetrics)) and dupRadar (v1.0.0)^25^. Differential expression analysis was performed on human-mapped counts derived from featureCounts using limma/voom^26,27^. Unless otherwise stated, we applied an absolute log2 fold-change cut-off of 1 and a false discovery rate (FDR) of <0.05.

Reported gene expression values for these QuantSeq RNA samples are given as TPMs (Transcripts Per Million) or CPMs (Counts Per Million).

### Principal Component Analysis

Principal component analysis of the RNA sequencing samples was performed using the plotPCA function implemented in the DESeq2 package in R^28^.

### Gene Clustering

Clustering of RNA sequencing samples was carried out using the pheatmap function from the pheatmap package in R. Clustering was based on row-wise z-scored CPM values per gene across conditions.

### EnrichR Analyses

Differentially expressed genes for each condition were submitted to the enrichR webserver (https://maayanlab.cloud/Enrichr/) and Gene Ontology (GO) term enrichment results were downloaded in CSV format^29^.

## Results

CRISPR depletion screen data across hundreds of cancer cell lines^30,31^ revealed that most BRAF^V600E^ mutant cell lines are dependent on DUSP4 (Figure 1A). Among 1,096 cell lines, 17 were classified as DUSP4-dependent, defined by a Chronos score < −1 (Supplementary Figure 1A). Notably, 76% of these DUSP4-dependent cell lines harbor a BRAF^V600E^ mutation. Other DUSP family members are not required for survival or proliferation in BRAF^V600E^ mutant cells, highlighting DUSP4 as the most relevant DUSP paralog in this mutational context.

**Figure 1:**
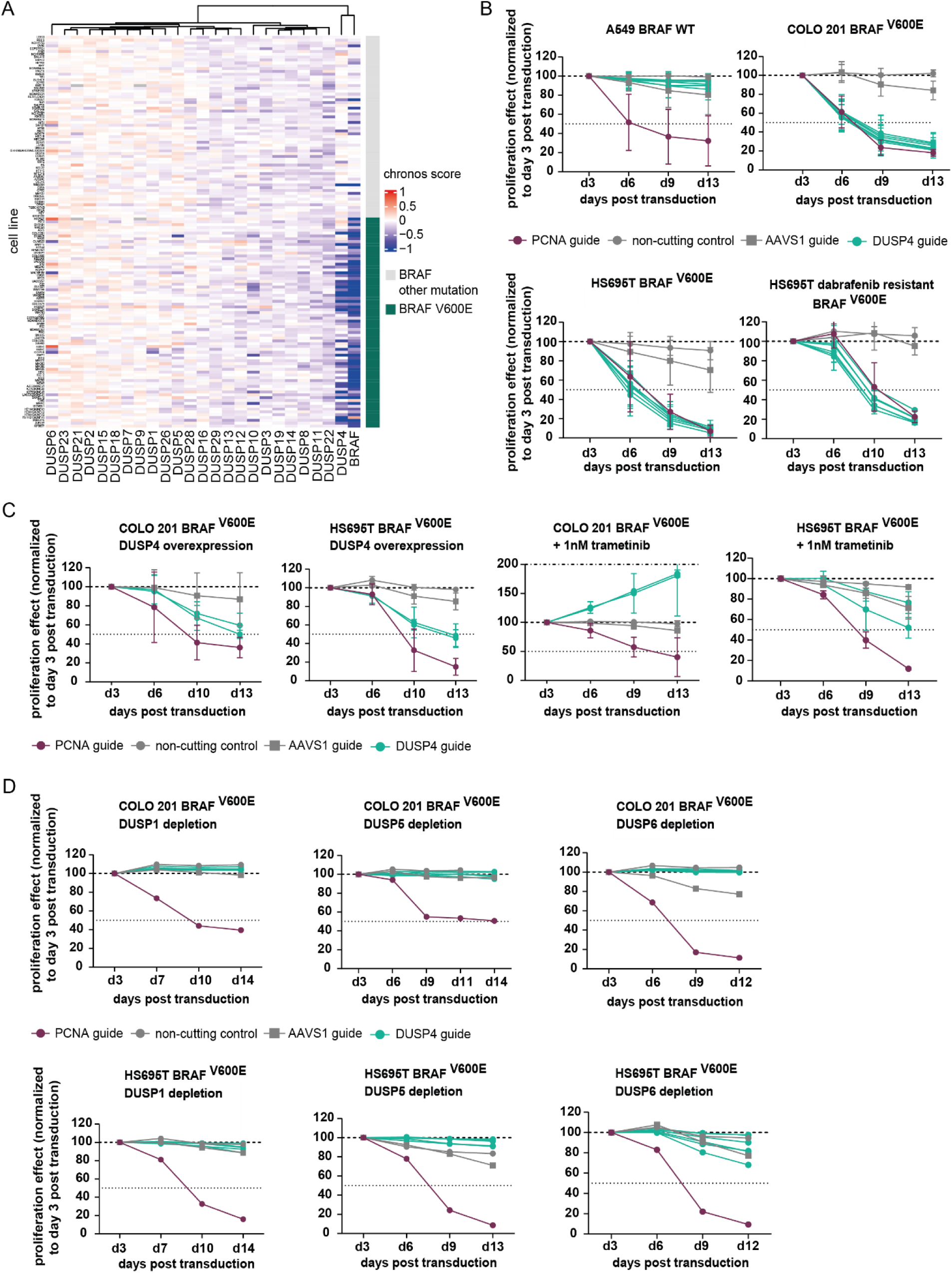
DUSP4 is a selective dependency in BRAF^V600E^ mutant cancer cell lines. **A,** DUSP4- and BRAF-dependent cell lines are enriched in the BRAF^V600E^ mutant cluster. Gene dependency data of the DUSP paralog family and BRAF originates from the DepMap Consortium database. Columns represent the genes; rows represent the cell lines; dependency Chronos scores are depicted in the tiles, ranging from red (no dependency) to blue (dependency). The dark turquoise annotation bar indicates the BRAF^V600E^mutant cell line; the grey annotation bar for BRAF non-V600E mutation carrying cell lines. **B,** Proliferation effects after DUSP4 CRISPR depletion in BRAF^V600E^ mutant COLO 201 and HS695T cell lines and in BRAF wild-type A549 cell line measured as GFP signal normalized to day 3 post DUSP4 CRISPR guide transduction. 2 biological replicates are shown. Strong proliferation defects are detected in BRAF^V600E^ mutant cell lines only. BRAF^V600E^ cell lines remain DUSP4 dependent in a BRAF inhibitor resistant setting. **C,** Proliferation effect after DUSP4 CRISPR depletion and DUSP4 overexpression or 1 nM trametinib treatment in BRAF^V600E^ mutant COLO 201 and HS695T cell lines. The proliferation effect is measured as GFP signal normalized to day 3 post DUSP4 CRISPR guide transduction. 2 biological replicates are shown. The DUSP4 proliferation effect can be partially rescued by DUSP4 overexpression and completely rescued by MAPK pathway inhibition upstream of DUSP4. **D,** Proliferation effect after CRISPR depletion of DUSP paralogs DUSP1, DUSP5 and DUSP6 measured as GFP signal normalized to day 3 post DUSP CRISPR guide transduction. DUSP1, DUSP5 or DUSP6 depletion does not cause a proliferation effect in COLO 201 and HS695T cell lines.

CRISPR depletion experiments in the BRAF^V600E^ mutant colorectal cancer cell line COLO 201 and the melanoma cell line HS695T confirmed the DUSP4 dependency of BRAF^V600E^ mutant cells (Figure 1B). DUSP4 protein levels were markedly reduced at day 4 post-transduction with DUSP4-targeting guide RNAs in both cell lines (Supplementary Figure 1B). Knockout of DUSP4 resulted in a pronounced reduction in proliferation in COLO 201 and HS695T cells. A similarly strong effect was observed in HS695T cells rendered resistant to the BRAF inhibitor dabrafenib. Conversely, no proliferation defects were observed upon DUSP4 loss of function in the BRAF wild-type cell line A549, confirming the BRAF^V600E^-specific nature of the DUSP4 dependency.

To confirm that the observed proliferation effects were on-target, we performed rescue experiments using doxycycline-inducible DUSP4 overexpression. Expression of the rescue construct (8-fold and 5-fold higher compared to parental cells) partially restored the phenotypes in both HS695T and COLO 201 cells (Figure 1C and Supplementary Figure 1C). Pharmacological inhibition of the MAPK pathway upstream of DUSP4 (by adding 1 nM of the MEK inhibitor trametinib to the medium) resulted in complete rescue and enhanced proliferation in COLO 201 cells. Similar effects were observed in HS695T cells (Figure 1C).

Depletion of closely related DUSP4 paralogs (DUSP1, DUSP5, and DUSP6) did not impair proliferation in COLO 201 and HS695T cell lines, suggesting a specific dependency on DUSP4 among the DUSP family members in the tested cell lines (Figure 1D and Supplementary Figures 1D and 1E).

To investigate the effect of DUSP4 knockdown *in vivo*, we generated colorectal BRAF^V600E^ mutant DUSP4 ARTi models (Figure 2A), in which an artificial RNA interference (ARTi) effector sequence was integrated into the C-terminus of DUSP4. The effector sequence, designed to mediate potent and off-target-free knock-down, enables highly stringent loss-of-function experiments in vitro and in vivo using doxycycline-inducible micro-RNA-embedded small hairpin RNA interference (shRNAmirs)^19,32–34^.

**Figure 2:**
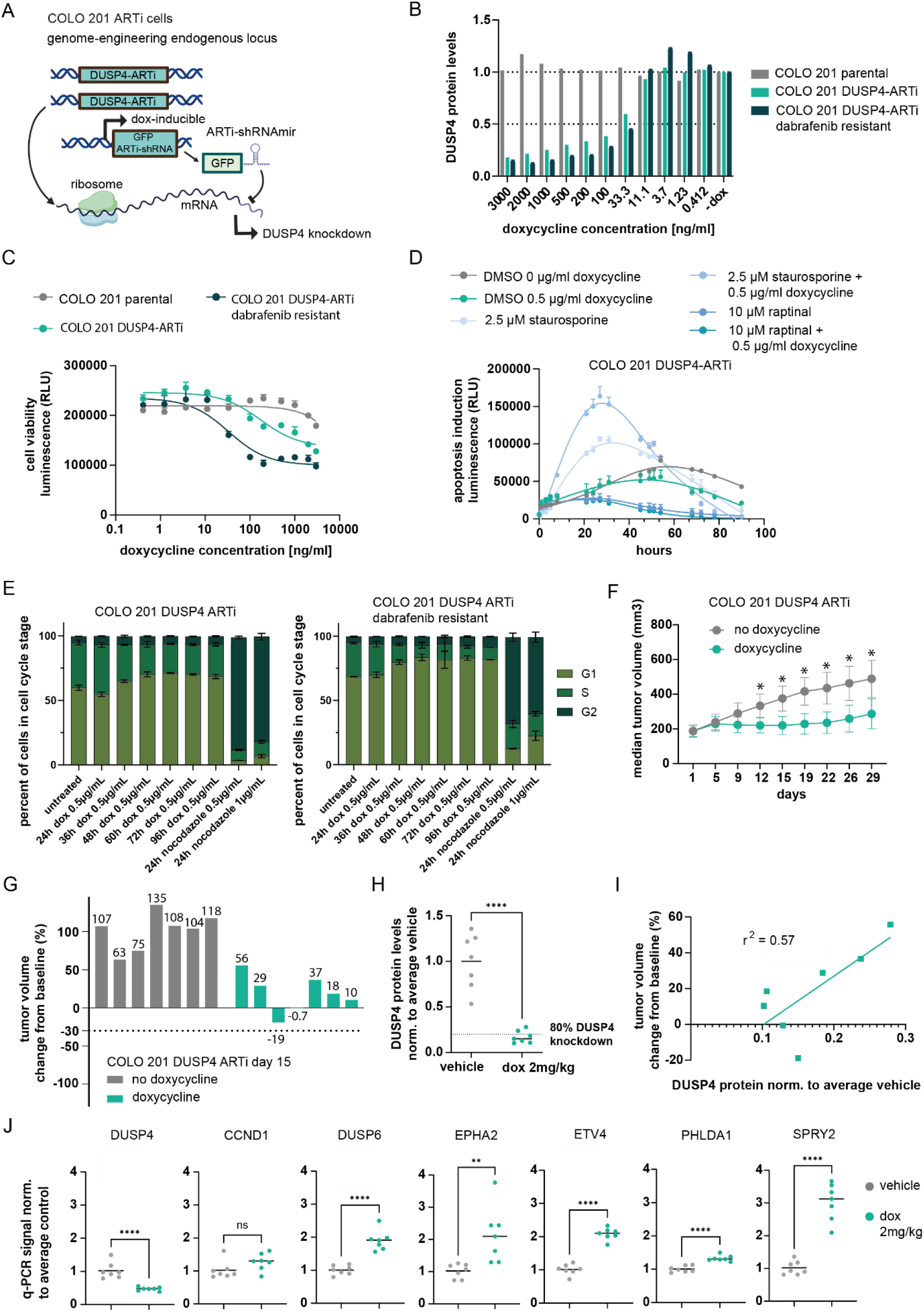
DUSP4 knockdown using the ARTi system leads to proliferation defects and G1 cell cycle arrest *in vitro* and *in vivo*. **A,** DUSP4 is C-terminally tagged with an ARTi sequence enabling RNAi interference and DUSP4 RNA degradation upon doxycycline mediated shRNAmir expression. **B,** DUSP4 protein levels after 6 days of doxycycline addition. A DUSP4 knockdown of approximately 50% is achieved at a concentration of 100 ng/ml doxycycline in untreated and dabrafenib resistant DUSP4 ARTi cell lines. **C,** Proliferation effect measured in CellTiter-Glo assay in untreated and dabrafenib resistant DUSP4 ARTi cell lines upon 6-day doxycycline induced DUSP4 knockdown. Error bars represent the standard deviation of duplicate measurements**. D,** Apoptosis signal measured as RLU luminescence in COLO 201 DUSP4 ARTi cell line upon doxycycline, staurosporine or raptinal treatment over 100 hours. Error bars represent the standard deviation of triplicate measurements. **E,** EdU staining and subsequent Flow Cytometry Assay results to infer cell cycle stages in COLO 201 DUSP4 ARTi and in dabrafenib resistant COLO 201 DUSP4 ARTi cells upon doxycycline mediated DUSP4 knockdown at 24 h, 36 h, 48 h, 60 h, 72 h and 96 h. Error bars represent the standard deviation of triplicate measurements. **F,** Median tumor growth curves of the COLO 201 DUSP4 ARTi efficacy model following doxycycline**-**induced DUSP4 knockdown. Error bars represent the standard deviation of tumor volume per day and treatment groups. **G,** Relative tumor volumes of the COLO 201 ARTi xenograft models are indicated as percent change from baseline at day 15. **H,** DUSP4 protein levels quantified by WES experiments after doxycycline induced DUSP4 knockdown in COLO 201 DUSP4 ARTi CDX models. Significant knockdown of protein levels is observed *in vivo*. The dashed line indicates 80% DUSP4 knockdown compared to average vehicle control. **I,** Correlation between tumor volume change from baseline and DUSP4 protein levels normalized to the average vehicle control, assessed by linear regression. r^2^ indicates the effect size. **J,** MPAS score gene expression levels for DUSP4, CCND1, DUSP6, EPHA2, ETV4, PHLDA1 and SPRY2 across subcutaneous DUSP4 ARTi tumors 29 days post**-**start of doxycycline treatment. Gene expression was measured by qPCR in triplicate per sample and normalized to GAPDH and the average control.

DUSP4 protein levels were reduced by approximately 80% compared to parental control at a concentration of >200 ng/mL doxycycline in both the COLO 201 DUSP4 ARTi and BRAF inhibitor-resistant COLO 201 DUSP4 ARTi models (Figure 2B and Supplementary Figure 2A). Consistent with this level of protein reduction, a significant decrease in proliferation was observed upon DUSP4 knockdown in both models compared to control cell lines. The proliferation effect in the dabrafenib-resistant model (Supplementary Figure 2B) was stronger than in dabrafenib naïve cells (Figure 2C).

To assess the terminal phenotype *in vitro*, we performed a RealTime-Glo™ Annexin V Apoptosis assay, which quantifies apoptotic activity via a luminescence signal. Staurosporine and raptinal treatments served as positive controls for apoptosis and necrosis induction. Neither apoptosis (Figure 2D) nor necrosis (Supplementary Figure 2C) increased substantially upon doxycycline-mediated DUSP4 knockdown. Next, we examined cell cycle arrest using the Click-iT® Plus EdU Flow Cytometry Assay. Following doxycycline-mediated DUSP4 knockdown, we observed an increase in the proportion of cells in the G1 phase in both COLO 201 DUSP4 ARTi parental and dabrafenib-resistant cells and a decrease in S-phase cells (Figure 2E), suggesting that impaired DUSP4 function leads to cell cycle arrest.

Next, we evaluated the anti-tumor efficacy of DUSP4 knockdown in a xenograft model using the COLO 201 ARTi cell line *in vivo*. In subcutaneous COLO 201 tumor-bearing mice, DUSP4 knockdown achieved by administering 2 mg/kg doxycycline via drinking water resulted in tumor stasis and a statistically significant tumor growth inhibition (TGI) of 82.2% on day 29 compared to the vehicle control (Figure 2F). The strongest effect was observed on day 15, with a TGI of 97% and 1 of 7 tumors showing regression to −19% (Figures 2F and 2G). Signs of tumor regrowth appeared from day 22 onward (Figure 2F). Biomarker analysis of tumors extracted on day 29 revealed a significant reduction in DUSP4 protein (80%) and RNA (50%) levels in 5 of 7 doxycycline-treated COLO 201 DUSP4 ARTi mice compared to control (Figure 2H and Supplementary Figure 2D), consistent with *in vitro* findings. Tumor volume change from baseline correlated with DUSP4 protein levels (r^2^ _linear regression_ = 0.57). The two least responsive COLO 201 DUSP4 ARTi CDX models showed the highest residual DUSP4 protein expression (Figure 2I).

All MPAS score genes were upregulated on day 29 of doxycycline treatment, except for DUSP4 (consistent with the shRNA mechanism of action) and cyclin D1 (CCND1) (Figure 2J and Supplementary Figure 2E). CCND1 encodes a regulatory subunit of the cyclin dependent kinases CDK4 and CDK6, whose activity is necessary for G1- to S-phase cell cycle transition. The unchanged CCND1 levels at day 29, may explain the onset of tumor regrowth despite continued doxycycline treatment and could indicate an adaptive response to DUSP4 loss of function.

To investigate the molecular changes associated with DUSP4 knockdown and the resulting proliferation defects, we performed RNA-seq following DUSP4 CRISPR knockout in COLO 201 and COLO 201 dabrafenib-resistant cells. Principal Component Analysis showed clustering of similarly treated replicates, as expected (Figure 3A). DUSP4 RNA levels were reduced after transduction with DUSP4-targeting CRISPR guides compared to AAVS1 control guides in both COLO 201 models (Figure 3B).

**Figure 3:**
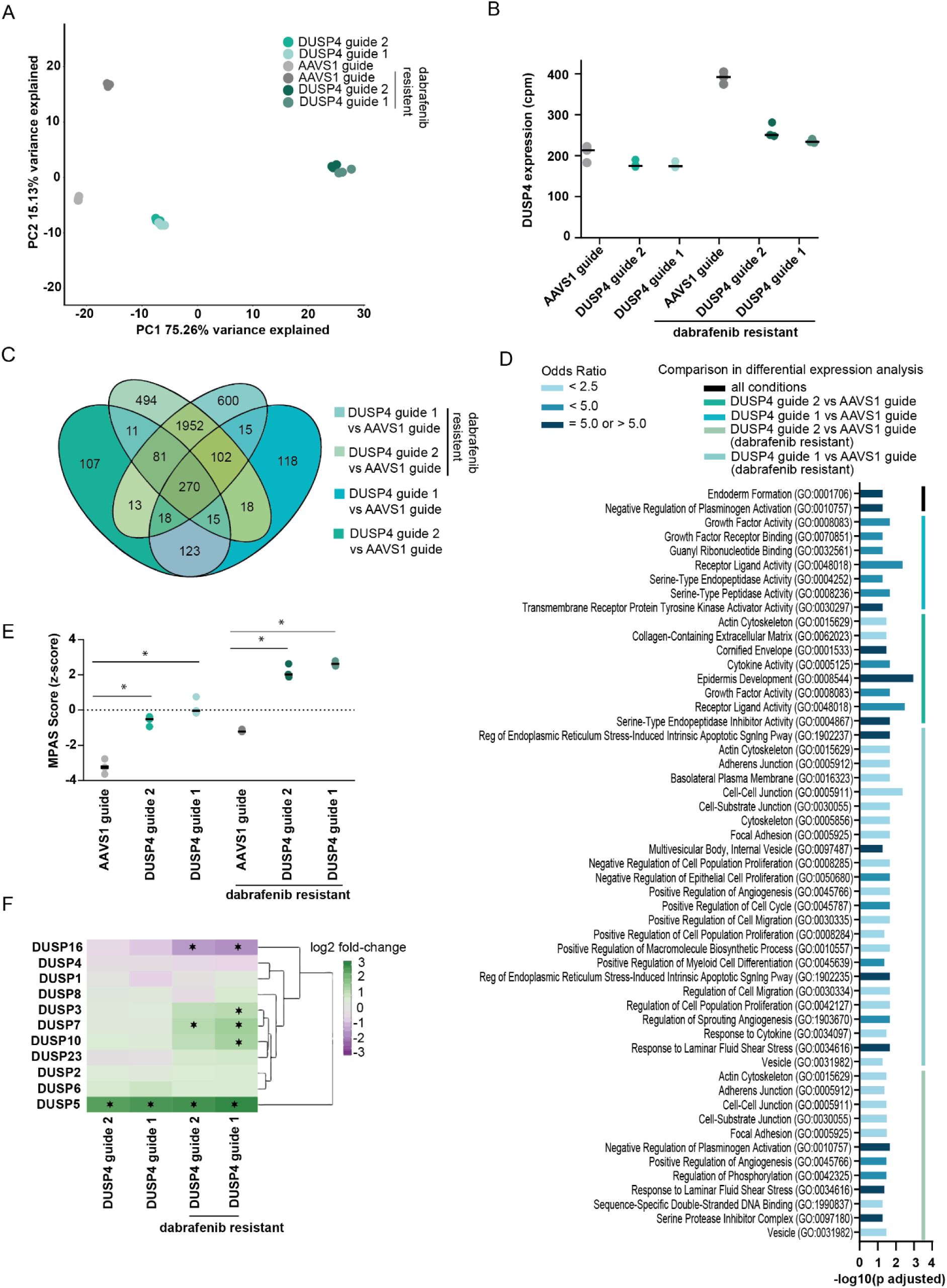
RNA sequencing reveals increased MAPK pathway activity and upregulated DUSP5 expression upon DUSP4 loss. **A,** Principal component analysis of RNA-sequencing data. The first two Principal Components and their variance explained are shown. **B,** DUSP4 RNA expression levels in counts per million mapped reads (CPM) measured in the RNA-seq experiment across COLO 201 cell line samples. **C,** Differentially expressed genes after DUSP4 CRISPR knockout compared to AAVS1 control knockout across conditions. Genes were considered significant at an absolute log2-fold change cut-off of 1 and a false discovery rate (FDR) < 0.05. **D,** GO term enrichment analyses for the differentially expressed genes per DUSP4 knockout condition. The y-axis shows GO term names, the x-axis shows the −log10 adjusted *p-value*. Enriched GO terms with an adjusted *p-value* of ≤ 0.05 are shown. The enrichment odds ratio is color coded in shades of blue. **E,** MPAS score depicted as z-score across CRISPR guide treatment conditions in COLO 201 wild-type and COLO 201 dabrafenib-resistant cell lines. Asterisks indicate significant changes (*p-value* < 0.05) when comparing DUSP4 guide treatment vs. AAVS1 control guide treatment. **F,** Significant differential expression of DUSP paralog family members upon DUSP4 CRISPR knockout compared to AAVS1 control knockout. Genes were considered significant at an absolute log2-fold change cut-off of 1 and an FDR < 0.05.

A total of 270 genes were differentially expressed (adjusted *p-value* < 0.05 and absolute fold change > 1) when comparing DUSP4 knockout vs. AAVS1 control across conditions (Figure 3C). GO enrichment analysis was performed using the enrichR webserver^29^ for each condition (Figure 3D). GO terms enriched at an adjusted p-value of 0.05 or lower are shown. Enrichment of terms related to endoderm and gut development was observed, likely reflecting the colorectal cancer lineage of COLO 201. Additionally, enrichment of GO terms associated with receptor protein tyrosine signaling and growth factor activity may be explained by increased MAPK pathway activity upon DUSP4 depletion. GO enrichment of serine protease inhibitor and plasminogen activity terms are driven by serine protease inhibitor 2 (SERPINE2) and urokinase-type plasminogen activator (PLAU) gene expression, both of which are linked to MAPK pathway activity^35,36^.

Relative MAPK pathway activity was assessed using the MPAS score, which is based on the expression levels of 10 MAPK pathway-activated genes and enables signal comparison across cell lines and under perturbations^37^. The MPAS score^37^ was significantly increased after DUSP4 knockout (Figure 3E), indicating that the MAPK pathway activity was markedly elevated in BRAF^V600E^ mutant COLO 201 cell lines upon DUSP4 loss. Higher MPAS scores were observed in dabrafenib-resistant cell lines under both control and DUSP4 loss-of-function conditions, suggesting that BRAF inhibitor-resistant cell lines require DUSP4 to dampen pathway activity even at elevated levels.

Changes in the expression of DUSP family members were observed following DUSP4 knockout; however, only DUSP5 expression was significantly altered across all conditions (Figure 3F).

To determine whether DUSP5 upregulation could serve as a compensatory mechanism for BRAF^V600E^ mutant cancer cells to mitigate DUSP4 loss, we quantified DUSP5 protein levels at 24, 48, 72 and 96 hours after doxycycline-induced DUSP4 knockdown using COLO 201 DUSP4 ARTi cell line models. DUSP5 levels increased consistently upon DUSP4 knockdown in both COLO 201 and a COLO 201 dabrafenib-resistant DUSP4 ARTi models compared to untreated controls at all time-points *in vitro* (Figure 4A). This observation was confirmed *in vivo*, where significant DUSP5 induction was observed in tumors on day 29 following doxycycline-induced DUSP4 knockdown (Figure 4B).

**Figure 4:**
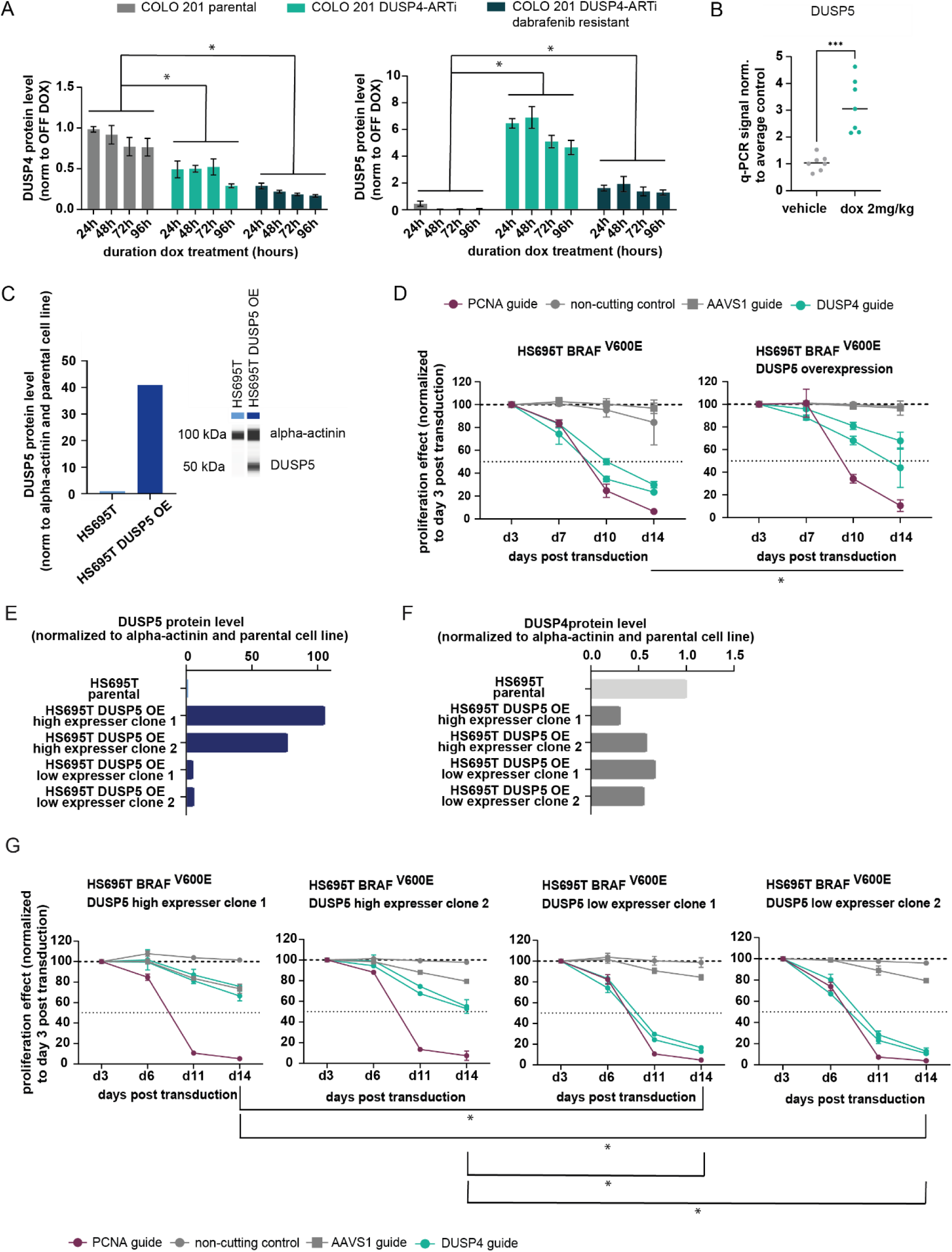
DUSP5 expression as a potential paralog compensating for DUSP4 loss. **A,** Quantification of DUSP4 and DUSP5 protein levels in COLO 201 parental, COLO 201 DUSP4-ARTi and COLO 201 DUSP4-ARTi dabrafenib resistant cell lines after 24, 48, 72 and 96 hours of doxycycline-induced DUSP4 knockdown normalized to doxycycline-untreated control. Asterisks indicate significant changes (*p-value* < 0.05) between conditions and individual time-points. **B,** DUSP5 RNA expression values in subcutaneous DUSP4 ARTi CDX tumors 29 days post start of doxycycline treatment. Gene expression was measured by qPCR in triplicate per sample and normalized to GAPDH and average control. **C,** Quantification of DUSP5 overexpression levels in HS695T cell line pool assessed by Simple Western^TM^ capillary based protein expression analysis. Protein expression levels are normalized to alpha-actinin and parental cell line levels. **D,** Proliferation after DUSP4 CRISPR depletion and DUSP5 overexpression cell line pool measured as GFP signal normalized to day 3 post DUSP4 CRISPR guide transduction. DUSP5 overexpression partially rescues the DUSP4 depletion phenotype in HS695T cells. Asterisks indicate significant changes (*p-value* < 0.05) between the average DUSP4 guide depletion effects per cell line condition at day 14 according to the unpaired t-test. **E,** Quantification of DUSP5 overexpression levels in HS695T DUSP5 high and low expressing clones assessed by Simple Western^TM^ capillary based protein expression analysis. Protein expression levels are normalized to alpha-actinin and parental cell line levels. **F,** Quantification of DUSP4 overexpression levels in HS695T DUSP5 high and low expressing clones assessed by Simple Western^TM^ capillary based protein expression analysis. Protein expression levels are normalized to alpha-actinin and parental cell line levels. **G,** Proliferation effect after DUSP4 CRISPR depletion and DUSP5 overexpression in DUSP5 high and low expressing clones measured as GFP signal normalized to day 3 post DUSP4 CRISPR guide transduction. Higher levels of DUSP5 overexpression rescue significantly stronger compared to low levels of DUSP5 overexpression when DUSP4 is depleted in HS695T cells. Asterisks indicate significant changes (*p-value* < 0.05) between the average DUSP4 guide depletion effects per cell line condition at day 14 according to the unpaired t-test.

We next overexpressed DUSP5 in the BRAF^V600E^ mutant melanoma cell line HS695T (Figure 4C) and performed DUSP4 CRISPR depletion assays to assess the impact on proliferation (Figure 4D). DUSP5 overexpression partially rescued the depletion effect caused by DUSP4 CRISPR depletion. The significant difference *(p-value* = 0.02) in the DUSP4 depletion effect between DUSP5 wild-type and DUSP5-overexpressing HS695T cells at day 14 suggests a potential functional redundancy between DUSP4 and DUSP5.

To further investigate the relationship between DUSP4 and DUSP5, we explored the extent to which DUSP5 overexpression could compensate for the loss of DUSP4 in BRAF^V600E^ mutant cells. Our initial experiments using a pooled cell population with varying levels of DUSP5 expression indicated a partial rescue of the DUSP4 phenotype. This observation led to two possible interpretations. First, it is possible that the functional redundancy between DUSP4 and DUSP5 is inherently incomplete, such that DUSP5 is unable to fully substitute for DUSP4 in HS695T cells, regardless of its expression level. Alternatively, the partial rescue could be attributed to heterogeneous DUSP5 expression within the pool, where only sub-clones with insufficient DUSP5 expression fail to fully compensate for DUSP4 loss, resulting in an intermediate phenotype across the population.

To distinguish between these possibilities, individual HS695T cell clones with different levels of DUSP5 expression (Figure 4E) were isolated and analyzed for DUSP4 expression levels (Figure 4F) and DUSP4 dependency (Figure 4G). The results supported the second hypothesis: clones exhibiting high DUSP5 expression demonstrated an almost complete rescue of the DUSP4 loss, whereas clones with low DUSP5 expression were unable to restore proliferation. In addition, DUSP4 levels decreased upon DUSP5 overexpression. These findings provide direct evidence that DUSP5 and DUSP4 are functionally redundant and are members of a bi-directional regulatory loop. Furthermore, they suggest that upregulation of DUSP5 is an adaptive mechanism employed by HS695T cells to compensate for the loss of DUSP4.

## Discussion

Oncogenic signaling is a key hallmark of human malignancies. This and previous studies provide evidence that a narrow gradient of MAPK signaling drives sustained proliferation and survival of cancer cells^13,16,38,39^. While inhibition of MAPK signaling leads to loss of proliferation and induction of apoptosis in MAPK-dependent tumor cells^7,10,20,40^, we show that overactivation of MAPK signaling through DUSP4 loss of function is associated with cell cycle arrest and reduced proliferative capacity in BRAF^V600E^ mutant cancer cells. These findings align with previous reports suggesting that MAPK pathway overactivation could be therapeutically exploited in BRAF^V600E^ mutant cells^14,17,18^.

To evaluate this therapeutic potential, we present the first *in vivo* study of DUSP4 knockdown in a BRAF^V600E^ colorectal cancer model. We demonstrate that DUSP4 loss of function results in tumor stasis *in vivo,* indicating potential disease control. However, MAPK overactivation via DUSP4 inhibition may not drive tumors into regression. These results suggest limited efficacy of DUSP4 inhibition in monotherapy.

Furthermore, our experiments provide evidence of a rapid adaptive response of cancer cells to DUSP4 inhibition. DUSP5 levels increase upon DUSP4 loss-of function, suggesting that tumor cells counteract elevated MAPK pathway activity by upregulating this paralog. While paralog redundancy amongst DUSPs has been documented previously, our data indicate that loss of one DUSP triggers rapid transcriptional re-programming in BRAF^V600E^ cancer cells to buffer MAPK signaling. This points to context-specific genetic buffering among DUSP proteins. Our study identifies DUSP5 as the primary paralog compensating for DUSP4 loss in colorectal cancer cells. DUSP5 is closely related to DUSP4, sharing high degree of sequence similarity, similar cellular localization, and comparable ERK1/2 affinity^41^. Consistent with our findings, DUSP5 has previously been associated with suppression of MAPK pathway activity in KRAS^G12D^ and HRAS^Q61L^ mutant cancers^42,43^.

Whereas the therapeutic impact of inhibiting oncogenic signaling is well established, the effects of hyperactivation only begin to emerge. Recent reports indicate that hyperactivation of oncogenic signaling may be incompatible with tumor cell survival^13,14,16^. In line with this concept, DUSP4 has been proposed as a cancer target for NRAS- and BRAF-mutant melanoma, particularly in the context of high Microphthalmia-associated transcription factor (MITF) expression^18^.

While CRISPR typically results in amorphic loss of function, RNAi more closely mimics the hypomorphic loss of function achieved with pharmacological inhibition^34^. Consistently, ARTi phenotypes closely resemble those observed with clinical-stage molecules, as demonstrated for the EGFR inhibitor osimertinib^19^ and the HER2 inhibitor zongertinib^44^. We therefore conclude that ARTi is a well-suited method to reliably assess the therapeutic potential of targets *in vivo* and approximate the outcome of pharmacological inhibition when inhibitors and/or degraders are not available. ARTi-mediated DUSP4 loss-of-function does not result in pronounced tumor shrinkage as observed for MEK or BRAF inhibition, precision oncology therapeutic targets in the same indication. While the therapeutic impact in humans can only be established in the clinic, our pre-clinical experiments suggest that DUSP4 inhibition i) will likely not achieve tumor shrinkage as monotherapy, ii) induces short-term pathway over-activation compensated by a DUSP5 regulatory loop and iii) cannot be combined with targeted therapy approved for BRAF^V600E^ mutant tumors, as MEK inhibition causes induction of cancer cell growth in the context of DUSP4 loss.

Collectively, we show that DUSP4 loss does not lead to tumor regression *in vivo* and provide evidence for a rapid adaptive response through upregulation of DUSP5. We therefore propose that DUSP4 inhibition as a monotherapy has limited therapeutic impact in BRAF^V600E^ mutant colorectal cancer.

## Data availability

The workflow for RNA-seq bioinformatics analyses is described in the Materials and Methods section. RNA sequencing data were uploaded to GEO under the accession number GSE289812 and will be made publicly available upon acceptance of this manuscript. All ARTi cell lines described in this study are available upon request.

## Acknowledgements

We thank Anna Obenauf, Johannes Zuber, and our collaborators at the IMP for scientific discussions and for providing the plasmids. We also thank Dominik Arnold, Sarah Oberndorfer, Alexandra Hörmann, Astrid Jeschko, Malgorzata Kamuda, and Julie Livingston for excellent technical assistance.

## Author contributions

SG and RAN designed the study, supervised the experiments, and wrote the manuscript. SG and DA performed bioinformatics analyses. BH and MK conducted the *in vitro* experiments and wrote the manuscript. MHH designed and supervised the *in vivo* experiments and reviewed the manuscript. YW provided feedback on the manuscript.

---

**Supplementary table 1:**
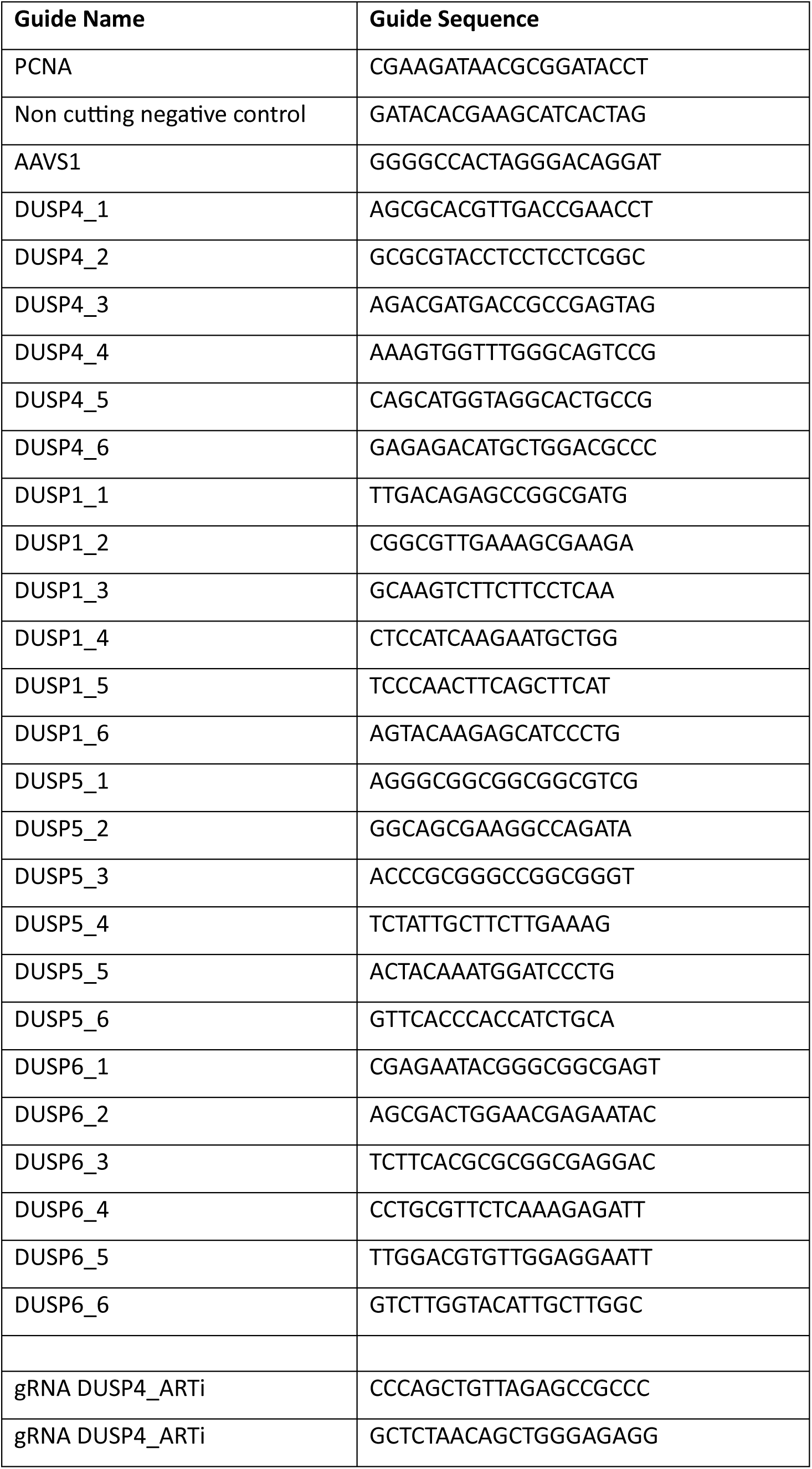
Guide sequences used in CRISPR depletion assays.

**Supplementary figure 1:**
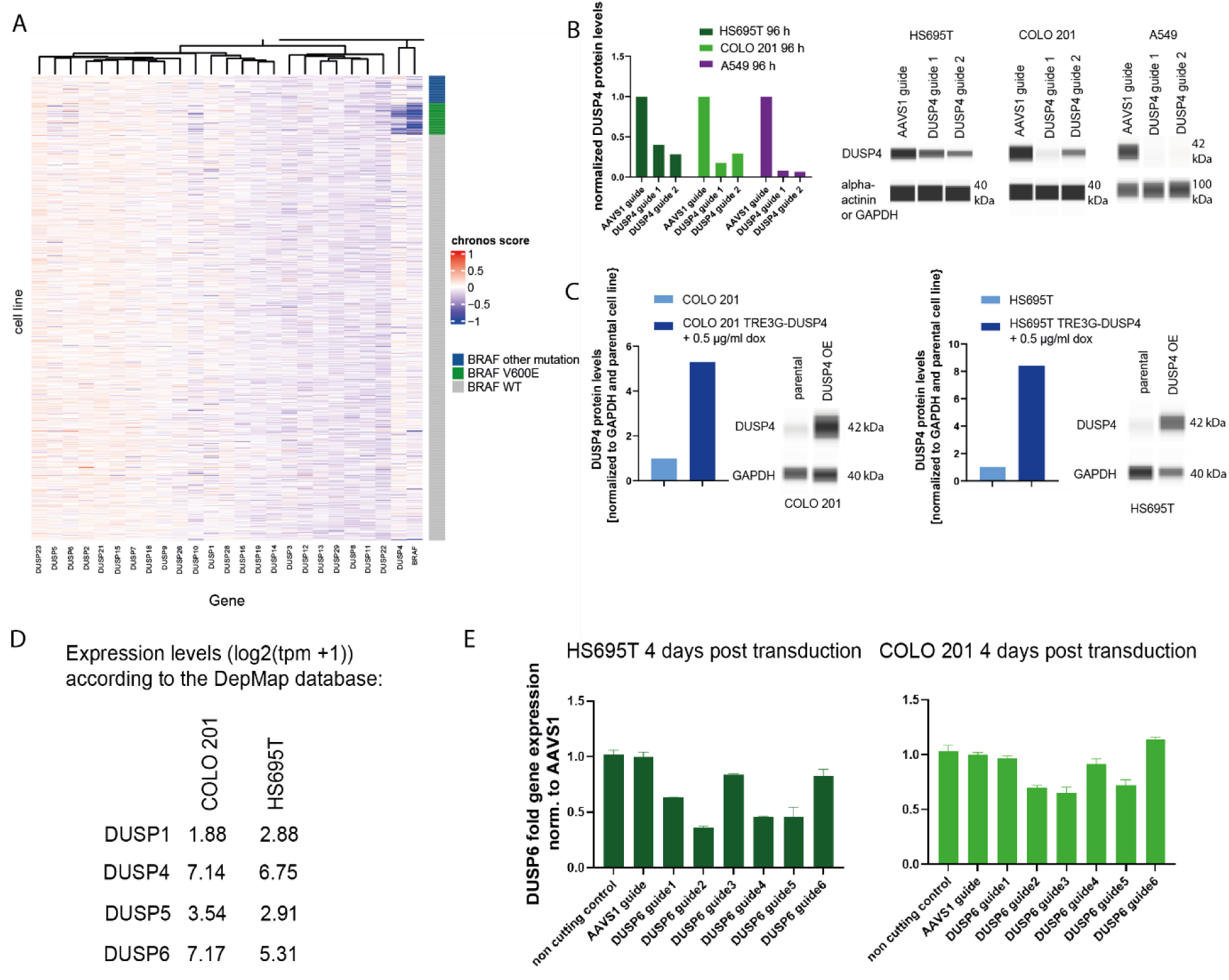
DUSP4 is a selective dependency in BRAF V600E mutant cancer cell lines. **A,** DUSP4 and BRAF dependent cell lines are enriched in the BRAF V600E mutant cluster. Gene dependency data for the DUSP paralog family and BRAF originates from the DepMap Consortium. Columns represent the genes; rows represent the cell lines; the dependency chronos score is depicted in the tiles from red (no dependency) to blue (dependency). The green annotation bar stands for BRAF V600E mutant cell lines, the dark blue annotation bar for cell lines carrying non-V600E BRAF mutations, and the grey annotation bar for BRAF wild-type cell lines. **B,** DUSP4 protein level analysis after DUSP4 CRISPR knockout at day 4 post-guide transduction in HS695T, COLO 201, and A549 cell lines using Western capillary electrophoresis (WES) quantification. Protein levels are normalized to GAPDH or alpha-actinin and DUSP4 levels after CRISPR depletion with AAVS1 control guide. **C,** WES quantification of DUSP4 protein overexpression levels after doxycycline addition in COLO 201 and HS695T cell lines. Protein levels were normalized to GAPDH and parental control cell line DUSP4 levels. **D,** DUSP1, DUSP4, DUSP5 and DUSP6 expression levels quantified as log2(tpm +1) in COLO 201 and HS695T cell lines according to RNA-sequencing data from the DepMap Consortium database. **E,** q-PCR quantification of DUSP6 levels after DUSP6 CRISPR depletion in COLO 201 and HS695T cell lines. RNA levels were normalized to GAPDH and AAVS1 guide treatment. Error bars depict the standard deviation of the triplicate measurements.

**Supplementary figure 2:**
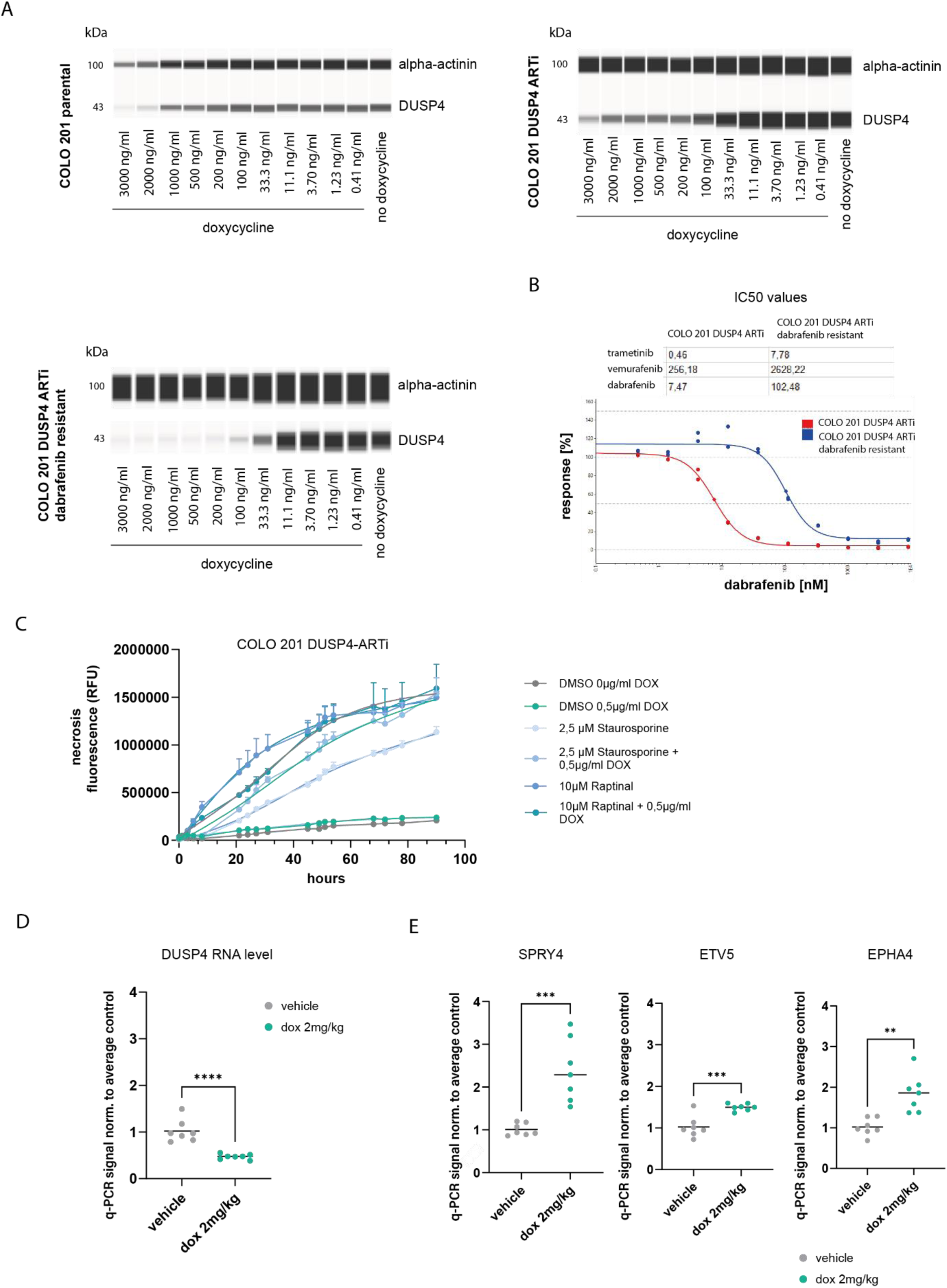
DUSP4 knockdown using the ARTi system leads to proliferation effects and G1 cell cycle arrest *in vitro* and *in vivo*. **A,** Images of DUSP4 knockdown levels induced by doxycycline addition in COLO 201, COLO 201 DUSP4 ARTi, and dabrafenib resistant COLO 201 DUSP4 ARTi cell lines generated by Western analysis using capillary electrophoresis (WES). **B,** Proliferation effect measured as relative IC_50_ value of COLO 201 DUSP4 ARTi and COLO 201 DUSP4 ARTi dabrafenib resistant cell lines upon trametinib, vemurafenib, or dabrafenib treatment. The dabrafenib resistant COLO 201 DUSP4 ARTi cell line shows an IC_50_ shift upon dabrafenib treatment compared to the compound naïve COLO 201 DUSP4 ARTi cell line. **C,** Necrosis signal measured as RFU fluorescence in COLO 201 DUSP4 ARTi cell line upon doxycycline, staurosporine, or raptinal treatment. No necrosis signal increase upon doxycycline mediated DUSP4 knockdown is observed compared to the necrosis inducing agent raptinal. **D,** DUSP4 RNA levels quantified by qPCR after doxycycline induced DUSP4 knockdown in COLO 201 DUSP4 ARTi CDX models. Significant knockdown of RNA levels are observed *in vivo*. **E,** MPAS score gene expression levels for SPRY4, ETV5 and EPHA4 across subcutaneous DUSP4 ARTi tumors 29 days post start of doxycycline treatment. Gene expression was measured by q-PCR in triplicates per sample and normalized to GAPDH and average control.

